# Expression of soluble Type IV Minor Pilins and isolation of a *Neisseria gonorrhoeae* PilI-PilJ subcomplex

**DOI:** 10.64898/2026.02.09.704877

**Authors:** Justin N. Applegate, Emma Y.D. Miller, Zoey R. Litt, Alfredo Ruiz-Rivera, Angela J.L. Lisovsky, Beth Traxler, Alexey J. Merz

## Abstract

Type IV pili and type II secretion systems assemble dynamic fibers used by bacteria and archaea for diverse functions. The pilus fiber is made up of major and minor pilin subunits containing a hydrophobic α-helical spine and a globular head. Purifying minor pilins is complicated by the hydrophobic α-helical spine, frequently present disulfide bonds, and low abundance within the fiber. These challenges have impeded structural and functional studies of pilin protomers. Here, we describe a method for expression and purification of soluble type IV pilin proteins from *Escherichia coli*. Signal peptidase I cleavage sites are engineered into the α-helix of the pilin proteins. This allows their globular domains to be purified from the periplasmic fraction. We used this method to obtain the *Neisseria gonorrhoeae* minor pilins PilI and PilK in soluble form. In a third case, where the minor pilin PilJ could not be obtained on its own, coexpression with PilI and purification of a PilI-PilJ heterodimer was possible. We suggest that PilI and PilJ form an obligate heterodimer that is essential for their function.

**Importance:** Type IV pili are essential to many bacteria responsible for disease. They can be found in both Gram-negative and Gram-positive bacteria, as well as archaea, making them likely present in the last common ancestor of all life on Earth. Despite their significance in a variety of species, there are large gaps in our understanding of the structure of these diverse biological machines. One roadblock to this research has been the difficulty of purifying the minor pilin proteins that serve different functions in the fiber. Here, we describe a novel method for the purification of these proteins and demonstrate the ability of this method to identify a protein-protein interaction between two minor pilins of *Nesseria gonorrhoeae*.

## Introduction

Type IV pili (T4P) are found in diverse organisms including Gram-negative and Gram-positive bacteria, as well as archaea. They were likely present in the last common ancestor of all life on Earth (Makarova et al. 2016). These protein fibers are used in twitching motility, surface sensing, biofilm formation, DNA uptake, and adhesion to host cells (Craig et al. 2019; Singh et al. 2022). T4P are essential virulence factors for many pathogenic organisms including *Neisseria gonorrhoeae*, the etiologic agent of gonorrhea. The pilus fiber is composed of pilin subunits, which include the major pilin that makes up most of the fiber, and minor pilins, which structurally resemble the major pilin but are present in the fiber in lower proportions. These minor pilins can serve several functions. In *N. gonorrhoeae*, PilV and ComP are involved in host cell adhesion and DNA uptake (Hughes-Games et al. 2022), while other pilins inlcuding PilI, PilJ, and PilK are essential for efficient pilus formation (Winther-Larsen et al. 2005).

Both major and minor prepilins consist of an N-terminal pilin signal sequence, a hydrophobic α-helical domain and a C-terminal globular head domain. The cytoplasmic N-terminal portion of the prepilin signal sequence is cleaved by a bifunctional enzyme, prepilin peptidase. Mature pilins are then N-methylated by the same enzyme (Nunn and Lory 1991). We follow the convention of numbering pilin amino acids from the first residue of the cleaved, mature pilin. The α-helical spines of pilins are embedded within the inner membrane, until the pilins are assembled into the fiber by the pilus assembly machine. T4P can also retract, depolymerizing from the base and releasing pilin protomers back into the cytoplasmic membrane pool. When an excessive quantity of pilin is present in the membrane and fiber assembly is disrupted, mature major pilins can be cleaved at the 39/40 residue, creating a soluble form of the globular domain, known as S-pilin. The protease responsible for this cleavage has not been identified (Giltner et al. 2012; Haas et al. 1987). It is not known whether S-pilin forms of the minor pilin proteins can occur.

Major pilin proteins have been purified in quantity from native fibers using mechanical shearing, followed by cycles of fiber precipitation. Pilin monomers or dimers are then obtained by dissolving the fibers in detergent, and major pilins can be further purified using ion exchange and size exclusion chromatography (Brinton et al. 1978; Parge et al. 1990; Gonzalez Rivera and Forest 2017). Purifying minor pilins through this approach is impractical, as there are only small numbers of minor pilin molecules per fiber. In the case of tip-located minor pilins, there is likely only one molecule per fiber.

Any purification approach must consider the pilin protomer’s hydrophobic transmembrane α-helical spine. Moreover — and in contrast to the related pilins of type II secretion systems — most type IV pilins contain one or more disulfide bonds, which form only after translocation of the globular domain across the cytopasmic membrane into the periplasm (or, in Gram positive bacteria and Archea, the extracellular compartment). Recombinant purification is a useful strategy for obtaining larger quantities of pilin proteins. The full-length major pilin of *Acidithiobacillus thiooxidans* was purified from *Escherichia coli* using a thioredoxin fusion and detergents to maintain solubility (Páez-Pérez et al. 2025). For biochemical and structural applications, it is valuable to have forms of minor pilin proteins that remain soluble in the absence of lipids or detergents. For this purpose, truncated pilin globular domains are often expressed. Some soluble domains of pilins have been recombinantly expressed in the cytoplasm of engineered *E. coli* cells with aberrant redox potential (Karuppiah et al. 2016; Craig et al. 2003). To allow truncated pilins to fold in the periplasm, they have at times been expressed as fusions to a periplasmic protein like maltose-binding protein (Helaine et al. 2007), or bearing signal sequences from periplasmic proteins such as OmpA (Hazes et al. 2000) or DsbA (Ramboarina et al. 2004). The type II secretion system (TIISS) is closely related to the type IV pilus. Structural data for TIISS minor pseudopilins, homologous to the minor pilins of T4P, was obtained by expressing the globular domains cytosolically in *E. coli*. In some cases this required refolding from inclusion bodies and years of effort (Korotkov and Hol 2008; Yanez et al. 2008; Korotkov et al. 2012). In contrast to TIISS pseudopilins, most T4P minor pilins have one, two, or more disulfide bonds, making these targets even more challenging than TIISS pseudopilins.

To probe the assembly pathway of a Type II secretion pseudopilin, Francetic et al. (2006) introduced a signal peptidase I (SPI) cleavage site that would allow processing if the pseudopilin was translocated into the periplasm. They observed a soluble form of the globular domain, similar to naturally occurring major S-pilins in the T4P system. We took inspiration from this approach to express and purify soluble T4P minor pilins from the *E. coli* periplasm. This method allows the proteins to fold as integral membrane proteins with oxidized disulfide bonds in the periplasm. Soluble globular pilins can then be recovered from the periplasmic space without the use of detergents that could interrupt protein-protein interactions or interfere with downstream assays.

## Materials and Methods

### Media and Buffers

LB medium contained 10 g/L Bacto Tryptone, 5 g/L Bacto Yeast Extract, 5 g/L NaCl, 1 mM NaOH. ZYM-5052 was prepared as described (Studier 2005). Sucrose Shock Buffer contained 0.1 M Tris-acetate at pH 7.8, 20% (m/v) sucrose, and 1 mM EDTA. 0.5 mM MgSO_4_ was used for osmotic shock. 10x PBS stock was prepared with 80 g/L of NaCl, 2.0 g/L KCl, 14.4 g/L Na_2_HPO_4_, 2.4 g/L KH_2_PO_4_. PBST contained 1x PBS with 0.1% Tween-20. Wash Buffer A contained 1x PBS + 10 mM imidazole, pH 7.8. Wash Buffer B contained 1x PBS + 350 mM NaCl + 10mM imidazole, pH 7.8. SDS-PAGE buffer contained 25 mM Tris Base, 192 mM Glycine, 0.1% SDS. 4X Laemmli buffer contained 0.250 M Tris base, 8% SDS, 40% glycerol, 20% 2-mercapto-ethanol, and Bromophenol blue.

### Reagents

Most chemicals were obtained from Sigma-Aldrich. Phenylmethylsulfonyl fluoride was from Fisher Scientific (P/2805/44) and Ni-Sepharose High Performance resin was from Cytiva (#17526802).

### Design of cleavage sites

*N. gonorrhoeae* minor pilin sequences were redesigned to contain a Signal Peptidase I cleavage site within the α-helical region (typically around residue +25, where the first residue of the processed, mature full-length pilin in a native context is defined as +1). SignalP v. 5.0 (Almagro Armenteros et al. 2019) was used to predict the ability of these sites to be recognized. Sequences were altered until SignalP v. 5.0 gave a probability over 0.6. We found that a more recent, machine learning-based release of SignalP (v. 6.0) (Teufel et al. 2022) was not suitable for this purpose, because it dominantly recognizes native prepilin peptidase processing sites in pilin signal sequences, rather than the introduced SPI cleavage sites. (Note that prepilin peptidases are not present in the *E. coli* hosts that we employ for expression.) Predictive structural modeling of the engineered minor pilins was performed using Alphafold2 *via* CollabFold (Mirdita et al. 2022), or Alphafold3 *via* Google Alphafold Server (Abramson et al. 2024).

### Cloning

Expression vectors were prepared with standard methods. In brief, a geneblock (IDT, Inc.) was designed to encode the designed minor pilin with an introduced SPI cleavage site and a C-terminal TEV-His_6_ tag. These sequences were amplified by PCR and inserted into the pET22B-derived plasmid pHIS-Parallel1 (Sheffield et al. 1999) between the NdeI and NcoI cut sites, by the method of (Gibson et al. 2009). For the bicistronic vector containing *pilI* + *pilJ*_short_, the open reading frames were inserted between NdeI and BlgI. Sequences for all expression vectors used are presented in the Supplemental Materials outlined in Table S1.

### Culture growth and expression conditions

Expression vectors were transformed into competent *E. coli* BL21(DE3) cells. A single colony was picked and added to LB medium supplemented with 100 ug/ml ampicillin. This culture was grown overnight shaking at 200 rpm at 37° C. 5 mL of seed culture was added to a 2 L baffled flask containing 1 L ZYM-5052 (Studier 2005), supplemented with 100 µg/ml ampicillin, then grown at 37° C, shaking at 200 rpm. When cells reached an optical density at 600 nm of 1.0, they were moved to a 16° C incubator and shaken at 200 rpm. Cultures were grown for an additional 18 hours before being harvested by centrifuging at 4670 x g for 15 min and 4° C (Sorvall H6000A, RC-3B Refrigerated Centrifuge).

### Isolation of Periplasmic Fraction

Immediately after centrifugation, pelleted cells were chilled on ice for 10 minutes then gently resuspended in 10 mL of ice cold sucrose shock buffer per 10 g of cell pellet. The cell suspension was chilled on ice for 10 minutes before being centrifuged at 4670 x g for 15 min and 4° C (Sorvall H6000A, RC-3B Refrigerated Centrifuge). The supernatant was collected. To the collected supernatant, 10x PBS was added to a final concentration of 1x, and 200 mM phenylmethylsulfonyl fluoride (PMSF) was added from a 2-propanol stock to a final concentration of 1 mM. The pelleted cells were chilled on ice for 10 minutes then gently resuspended in 5 mL of ice-cold 0.5 mM MgSO_4_ per 10 g of cells. The cells were incubated on ice for 10 minutes in MgSO_4_ then centrifuged at 4670 x g for 15 minutes and 4 °C (Sorvall H6000A, RC-3B Refrigerated Centrifuge). The supernatant was collected. 10x PBS was added to a final concentration of 1x and 200 mM PMSF was added to a final concentration of 1 mM.

### Immobilized metal affinity purification

Supernatants from both the sucrose shock and 0.5 mM MgSO_4_ steps contained soluble minor pilins, so these fractions were pooled. For every 10 mL of pooled supernatant, 50 µL of packed Ni^2+^-NTA sepharose HP (Cytiva) was equilibrated in 1X PBS. Equilibrated resin was added to the pooled supernatant and bound in batch, nutating for 1 hr on ice. Samples were then centrifuged for 8 minutes at 500 x g and 4° C (TX-400 rotor, Sorvall Legend X1R). Supernatant was removed and resin was washed 3 times, with resuspension in wash buffer and centrifugation as in the previous step. For the first wash, the resin was resuspended in 5 mL of Wash Buffer A. For the second wash, the resin was resuspended in 10 mL of Wash Buffer B. For the third wash, the resin was resuspended in 5 mL Wash Buffer A. After washing the resin slurry was transferred to a chilled column. Bound material was eluted from the column using elution buffer (PBS + 350 mM imidazole), collected in 0.5 mL fractions.

### SDS-PAGE and immuno-blotting

Samples were mixed with 4x Laemmli buffer containing 20% 2-mercapto-ethanol (5% v/v final) then incubated at 95° C for 5 minutes. Samples were loaded into 12.5% (w/v) SDS-PAGE gels and run for 60 minutes at 190 V. Gels were either stained with Coomassie blue or for immuno blotting transferred to a 0.45 µm nitrocellulose membrane (Schleicher & Schuell BA85). Membranes were blocked with PBS Blocking Buffer (LICORbio) for 1 hr at room temperature then incubated with primary antibody (His.H8, Invitrogen MA1-21315) for 1 hour at room temperature. The membrane was rinsed in PBST 5 times for 5 minutes before being incubated with secondary antibody diluted 1:10,000 (Goat anti-Mouse IgG H+L Highly Cross-Adsorbed Secondary Antibody Alexa Fluor 488, Invitrogen A32723TR) for 45 minutes at RT. Membrane was then rinsed in PBST 5 times for 5 minutes and imaged using an Odyssey imaging system (LI-COR Biotech).

## Results and Discussion

### Design of SP1-celavable pilins

Type IV pilins are substrates of the PilD prepilin peptidase, which cleaves at the cytoplasmic face of the plasma membrane. Pilins do not natively contain SPI signal sequences, which are cleaved at the periplasmic face of the plasma membrane. SPI sequences are defined by a positively charged N-terminal region, a central hydrophobic region, a helix-breaking residue at position -4 to -6 relative to the cleavage site, and compact amino acid residues at positions -1 and -3 (Kaushik et al. 2022). The minor pilin PilK from *N. gonorrhoeae* strain MS11A was engineered to introduce a SPI cleavage site within its *α*-helical spine. This was accomplished by iterative cycles of *in silico* mutagenesis and signal sequence prediction, using the SignalP v. 5.0 algorithm (Almagro Armenteros et al. 2019). In PilK, we replaced the region from T37 to Y42 (TAAQSY) with PGAQASD (Figure 1B). SignalP v. 5.0 predicted a SPI cleavage site between A41 and S42 with a probability of 0.647. By contrast, in wild type PilK, the SPI cleavage probability was predicted to be 0.064 (Supplementary Figure 1).

**Figure 1.**
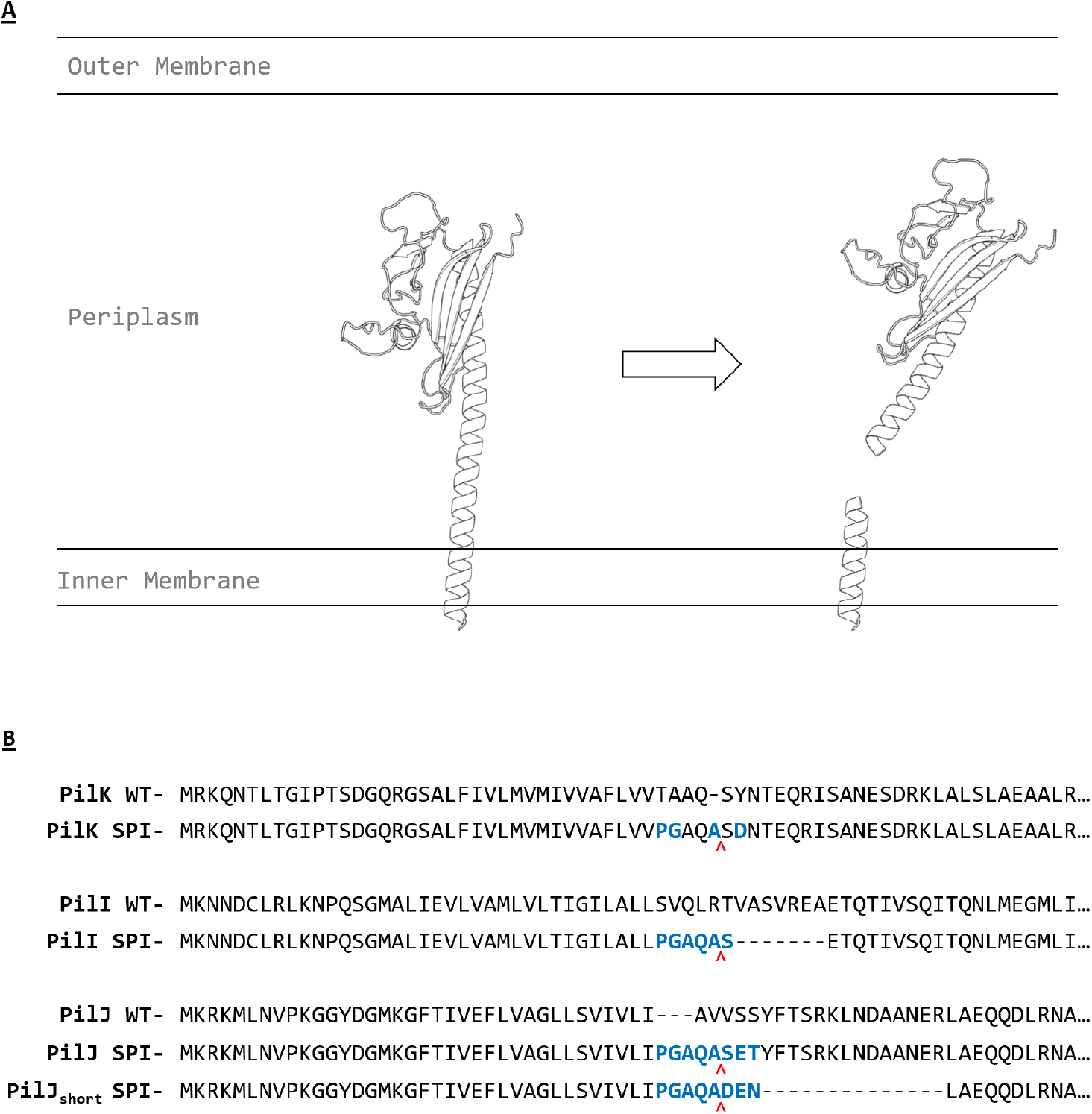
Signal Peptidase I cleavage of engineered minor pilins. **A**. A model for the post translational processing of pilins is shown. Signal Peptidase I cuts at an engineered SPI cut site. **B**. N-terminal sequences of minor pilins containing engineered Signal Peptidase I (SPI) are shown. Residues that diverge from the wild type are in blue. Red arrowheads indicate the engineered cleavage sites.

A plasmid was prepared to express the putatively cleavable variant of PilK in *E. coli*. Cultures were grown, expression was autoinduced, and a periplasmic fraction was isolated. In anti-His_6_ immunoblots, the whole-cell lysate contained two labeled species corresponding to the predicted masses of full-length and SPI-cleaved PilK-His_6_. In contrast, the soluble periplasmic fraction contained only the faster-migrating species, corresponding to SPI-cleaved, truncated PilK-His_6_ (Figure 2B). This indicates that the minor pilin is expressed and that the globular domain is processed at the engineered SPI cut site. Following immobilized metal chromatography, the soluble PilK-His_6_ fragment had a high yield of up to 20 mg of protein per L of starting culture (Figure 2A).

**Fig. 2.**
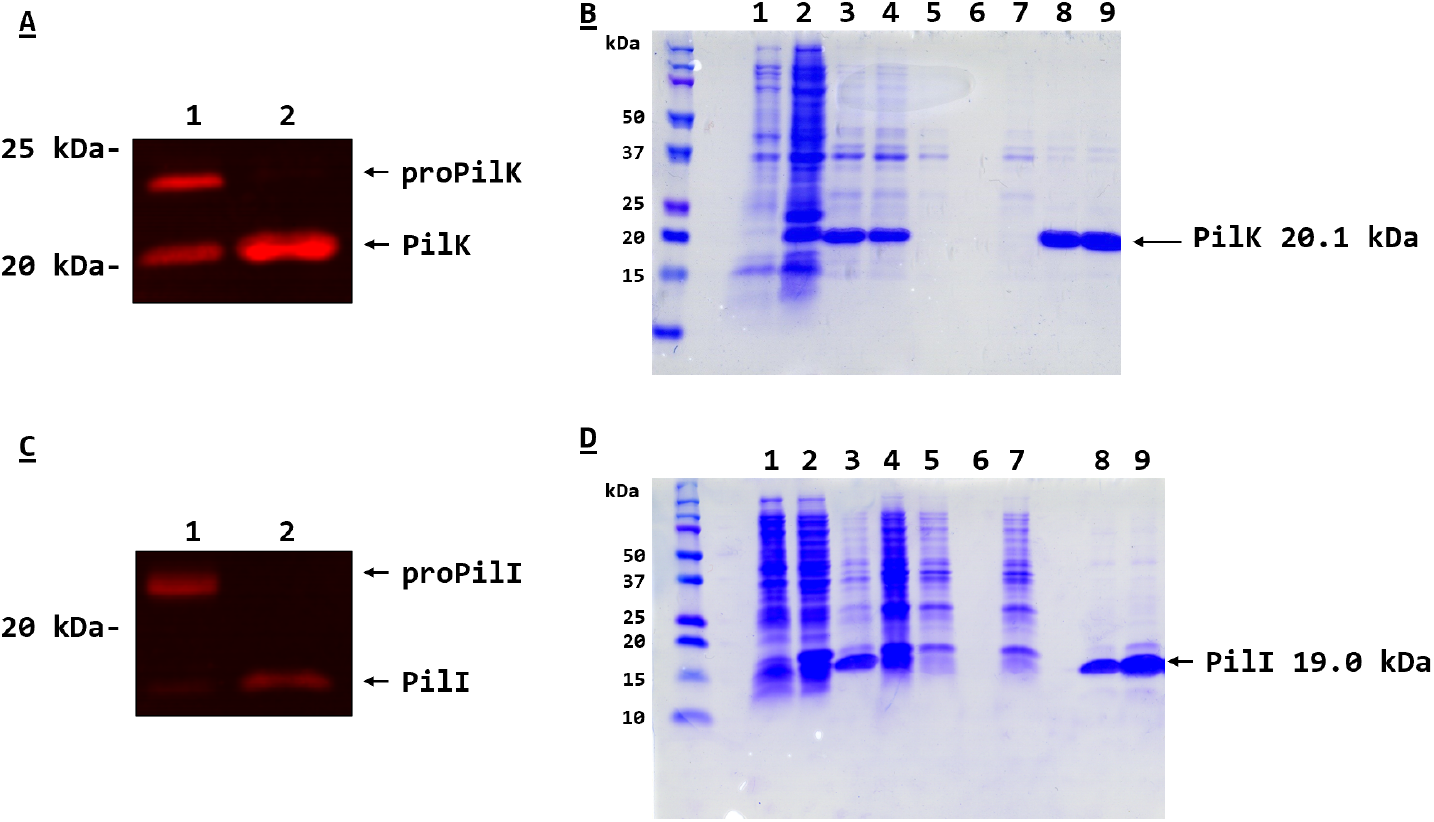
Introduced SPI sites yield soluble minor pilin fragments. **A**. Expression of engineered soluble minor pilin PilK-His_6_ was analyzed by anti-His_6_ immunoblot. Lane 1: whole cell lysate. Lane 2: periplasmic fraction. **B**. Purification of PilK-His_6_. Lane 1: pre-induction cell lysate. Lane 2: post-induction cell lysate. Lane 3: sucrose shock supernatant. Lane 4: MgSO_4_ supernatant. Lane 5: MgSO_4_ fraction affinity resin flowthrough. Lane 6: MgSO_4_ final wash. Lane 7: sucrose shock affinity resin flowthrough flowthrough. Lanes 8 and 9: eluted soluble PilK-His_6_. **C**. Expression of engineered minor pilin PilI-His_6_ was analyzed by anti-His_6_ immunoblot. Lane 1: whole cell lysate. Lane 2: periplasmic fraction. **D**. Purification of PilI-His_6_. Lane 1: pre-induction cell lysate. Lane 2: post-induction cell lysate. Lane 3: sucrose shock supernatant. Lane 4: MgSO_4_ supernatant. Lane 5: MgSO_4_ affinity matrix flowthrough. Lane 6: final affinity resin wash. Lane 7: sucrose shock affinity matrix flowthrough. Lanes 8 and 9: eluted soluble PilI-His_6_.

Putatively cleavable variants of PilI and PilJ were also designed (Figure 1B). For the engineered PilI-His_6_, SignalP5.0 predicted the likelihood of SPI cleavage to be 0.675, and for PilJ-His_6_, 0.565. PilI expressed well, with a yield of approximately 16 mg per L of culture (Figure 2D). Although a variety of growth and induction conditions were tested, we were unable to purify PilJ-His_6_ using the initial design (Figure 3A).

**Figure 3.**
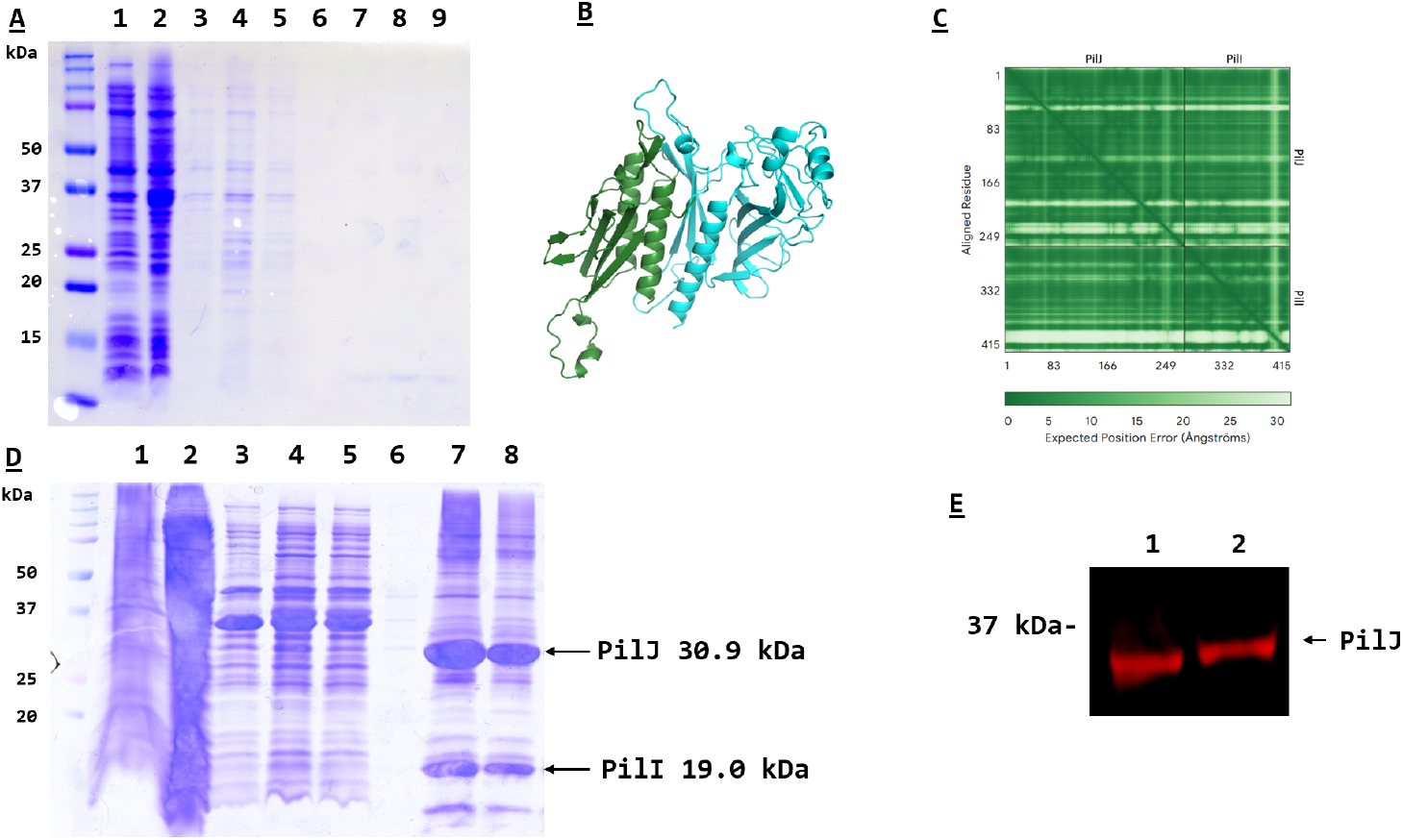
Expression and partial purification of a soluble PilI-PilJ complex. **A**. Expression and failed purification of soluble engineered PilJ-His_6_. Lane 1: pre-induction cell lysate. Lane 2: Post-induction cell lysate. Lane 3: sucrose shock supernatant. Lane 4: MgSO_4_ supernatant. Lane 5: affinity resin flowthrough of the pooled shock supernatants. Lane 6: final affinity resin wash. Lanes 7-9: affinity resin eluate fractions. **B**. Alphafold3 prediction of heterodimer formation between truncated PilI (Green) and PilJ (Cyan) soluble domains. **C**. Alphafold3 confidence matrix for the predicted soluble PilI-PilJ heterodimer. **D**. Purification of PilI PilJ_Short_-His_6_. Lane 1: pre-induction cell lysate. Lane 2: post-induction cell lysate. Lane 3: sucrose shock supernatant. Lane 4: MgSO_4_ shock supernatant. Lane 5: affinity resin flowthrough of the pooled shock supernatants. Lane 6: Final affinity resin wash. Lanes 7 and 8: Eluted PilJ_Short_-His_6_ complex. **E**. Expression of engineered soluble minor pilin PilJ_Short_-His_6_, co-expressed with engineered soluble PilI, was analyzed by anti-His_6_ immunoblot. Lane 1: whole cell lysate. Lane 2: periplasmic fraction.

To understand why the engineered PilJ-His_6_ may have expression issues, we used Alphafold3 to predict its structure. We found that compared to the minor pilins that expressed well, this design of PilJ had a larger hydrophobic region on the C-terminal side the SPI cleavage site. The cleavage site could not be moved downstream without interfering with its ability to be cut. Thus, another construct for PilJ expression, PilJ_short_-His_6_, was constructed, lacking 14 residues downstream of the cleavage site (Figure 1B). SignalP v. 5.0 predicted the likelihood of PilJ_short_-His_6_ SPI cleavage to be 0.582. Although PilJ_short_-His_6_ was expressed, we were still unable to purify it in soluble form.

### Purification of a stable PilI-PilJ subcomplex

In a previous study, deletion of the *pilJ* gene in *N. gonorrhoeae* resulted in decreased PilI expression at steady state. Moreover, deletion of the *pilI* gene in *N. gonorrhoeae* resulted in almost undetectable PilJ expression (Winther-Larsen et al. 2005). These results suggested the hypothesis that PilI and PilJ form a binary complex that stabilizes both proteins. In TIIS systems, the nearest paralogs of *N. gonorrhoeae* PilI and PilJ are Gsp/Eps I and J. Crystal structures of TIIS pseudopilins from *Vibro vulnificus* and enterotoxigenic *E. coli* revealed I-J dimers with extensive binding interfaces (Yanez et al. 2008; Korotkov and Hol 2008; Lam et al. 2009). Similarly, our Alphafold3 modeling of PilI-PilJ predicted with high confidence that PilI and PilJ form a stable heterodimer with the monomers in the same general arrangement as in the TIIS crystal structures (Figure 3B). Based on this, a bicistronic vector co-expressing both PilI and PilJ_short_ was prepared. Expression from this vector allowed for the purification of a heterodimer of PilJ_short_-His_6_ and PilI (Figure 3D). When run on an anti-His_6_ immunoblot, both whole cell lysate and periplasmic fractions contained only a single PilJ_short_-His_6_ band, with a mass of approximately 30 kDa, the size of the predicted soluble cleavage product (Figure 3E). A band corresponding to the uncleaved precursor was not detected. This may indicate that the SPI site is processed more efficiently than those of engineered PilK or PilI alone.

Our findings strongly support the idea that PilI association stabilizes PilJ, and suggests that the *Neisseria* T4P tip complex and TIIS pseudopili share a conserved quaternary subunit structure. In a companion study (Ellison *et al*., 2026 preprint), we present evidence that the *N. gonorrhoeae* PilI-PilJ heterodimer has a 1:1 stoichiometry, and that it functions as a recognition module for PilK and the tip adhesin PilC. In a working model, the PilI-PilJ module monitors formation of a complex between PilK and ∼10 residues at the extreme C-terminus of the tip-located adhesin PilC. The resulting Pil I-J-K-C heterotetramer would then license polymerization of the helical T4P fiber through a templating mechanism. Purification of minor pilins is essential for studying type IV pili. Recombinant expression from *E. coli* has been useful but has posed a variety of technical challenges. The complementary approach outlined here allows for expression and purification of these minor pilins in good yields, without the use of detergents that could interfere with protein-protein interactions. As we have seen with PilJ, interactions of these proteins can be essential to their stability and function. Further structural and biophysical characterization of minor pilin interactions is needed to understand the assembly, dynamics, and functions of type IV pili and type II secretion systems.

## Acknowledgments

J.N.A. was supported by the University of Washington STD & AIDS Research Training Program (NIH T32 AI07140). This work was supported by the National Institutes of Allergy and Infectious Disease (NIAID) R21 AI155991 and a kind gift to the UW Bacterial Research Fund from Professor J.M. Griffiss.

## Supplementary Tables

**Table S1.**
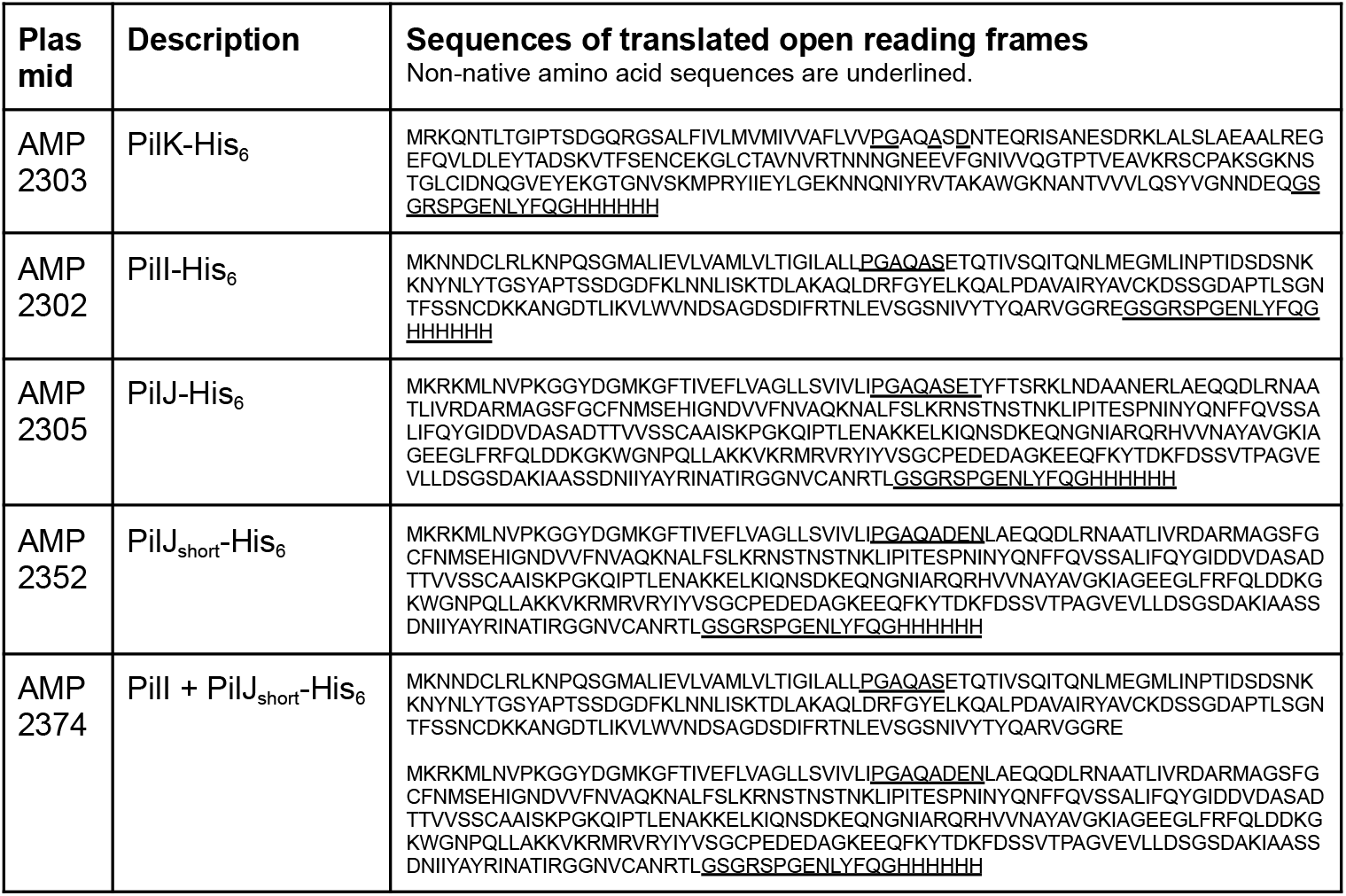
Plasmids used in this study.

**Table S2.**
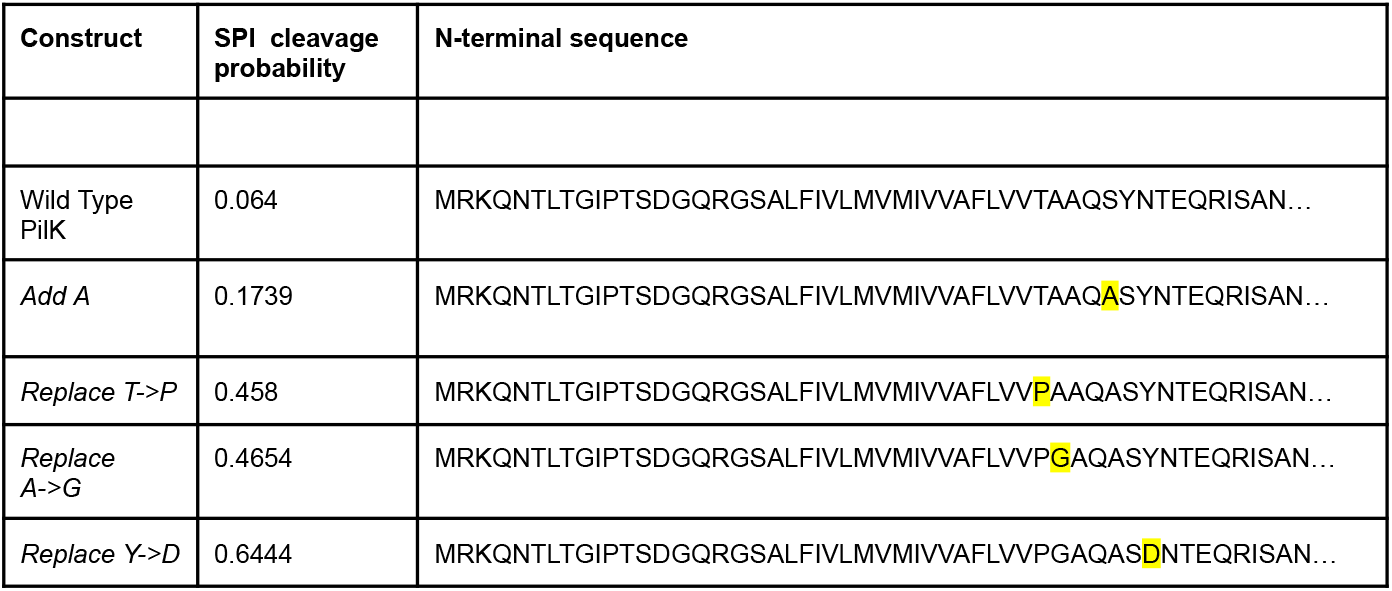
Iterative process for SPI site redesign of PilK. Changes made to the amino acid sequence in each iteration of design are highlighted in yellow. The predicted probabilities of SPI cleavage, estimated using SIGNALP5.0, are shown.

## Supplementary Figure

**Supplementary Figure S1.**
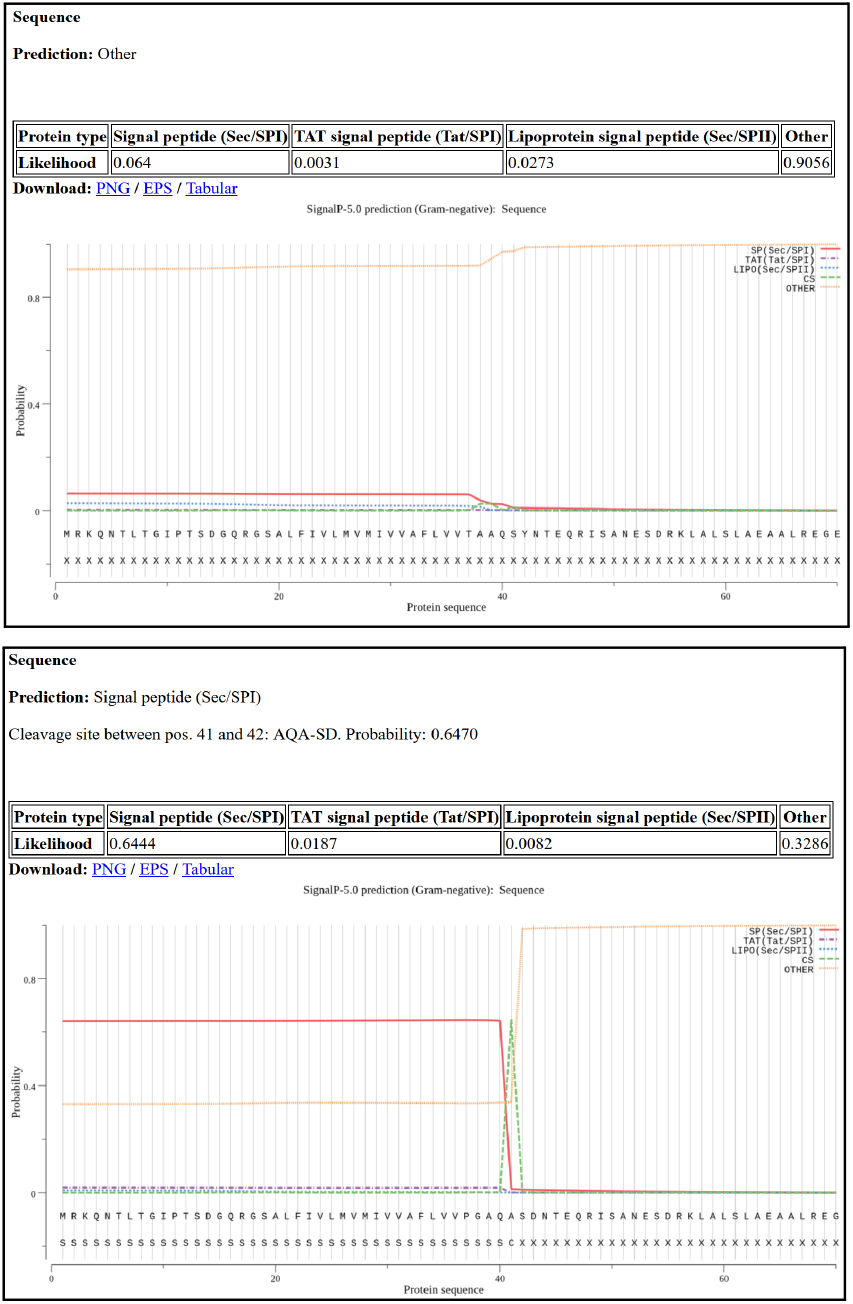
The top image shows the output of SignalP v. 5.0 for wild type PilK. The bottom image shows the output of SignalP v. 5.0 for the final iteration of the engineered PilK.

## Citations

Abramson, Josh, Jonas Adler, Jack Dunger, et al. 2024. “Accurate Structure Prediction of Biomolecular Interactions with AlphaFold 3.” Nature 630 (8016): 493–500. 10.1038/s41586-024-07487-w.

Almagro Armenteros, José Juan, Konstantinos D. Tsirigos, Casper Kaae Sønderby, et al. 2019. “SignalP 5.0 Improves Signal Peptide Predictions Using Deep Neural Networks.” Nature Biotechnology 37 (4): 420–23. 10.1038/s41587-019-0036-z.

Brinton, C., J. Bryan, J. Dillon, et al. 1978. “Uses of Pili in Gonorrhea Control Role of Bacterial Pili in Disease Purification and Properties of Gonococcal Pili and Progress in the Development of a Gonococcal Pilus Vaccine for Gonorrhea.” July 28. https://www.semanticscholar.org/paper/Uses-of-pili-in-gonorrhea-control-role-of-bacterial-Brinton-Bryan/ef7aba5ceb3bd9916d976cf9028d07ec68f8509e.

Craig, Lisa, Katrina T. Forest, and Berenike Maier. 2019. “Type IV Pili: Dynamics, Biophysics and Functional Consequences.” Nature Reviews. Microbiology 17 (7): 429–40. 10.1038/s41579-019-0195-4.

Craig, Lisa, Ronald K Taylor, Michael E Pique, et al. 2003. “Type IV Pilin Structure and Assembly: X-Ray and EM Analyses of Vibrio Cholerae Toxin-Coregulated Pilus and Pseudomonas Aeruginosa PAK Pilin.” Molecular Cell 11 (5): 1139–50. 10.1016/S1097-2765(03)00170-9.

Gibson, Daniel G., Lei Young, Ray-Yuan Chuang, J. Craig Venter, Clyde A. Hutchison, and Hamilton O. Smith. 2009. “Enzymatic Assembly of DNA Molecules up to Several Hundred Kilobases.” Nature Methods 6 (5): 5. 10.1038/nmeth.1318.

Giltner, Carmen L., Ylan Nguyen, and Lori L. Burrows. 2012. “Type IV Pilin Proteins: Versatile Molecular Modules.” Microbiology and Molecular Biology Reviews: MMBR 76 (4): 740–72. 10.1128/MMBR.00035-12.

Gonzalez Rivera, Alba Katiria, and Katrina T. Forest. 2017. “Shearing and Enrichment of Extracellular Type IV Pili.” Methods in Molecular Biology (Clifton, N.J.) 1615: 311–20. 10.1007/978-1-4939-7033-9_25.

Haas, R., H. Schwarz, and T. F. Meyer. 1987. “Release of Soluble Pilin Antigen Coupled with Gene Conversion in Neisseria Gonorrhoeae.” Proceedings of the National Academy of Sciences of the United States of America 84 (24): 9079–83. 10.1073/pnas.84.24.9079.

Hazes, Bart, Parimi A Sastry, Koto Hayakawa, Randy J Read, and Randall T Irvin. 2000. “Crystal Structure of Pseudomonas Aeruginosa PAK Pilin Suggests a Main-Chain-Dominated Mode of Receptor Binding1.” Journal of Molecular Biology 299 (4): 1005–17. 10.1006/jmbi.2000.3801.

Helaine, Sophie, David H. Dyer, Xavier Nassif, Vladimir Pelicic, and Katrina T. Forest. 2007. “3D Structure/Function Analysis of PilX Reveals How Minor Pilins Can Modulate the Virulence Properties of Type IV Pili.” Proceedings of the National Academy of Sciences of the United States of America 104 (40): 15888–93. 10.1073/pnas.0707581104.

Hughes-Games, Alex, Sean A. Davis, and Darryl J. Hill. 2022. “Direct Visualization of Sequence-Specific DNA Binding by Gonococcal Type IV Pili.” Microbiology (Reading, England) 168 (8). 10.1099/mic.0.001224.

Karuppiah, Vijaykumar, Angela Thistlethwaite, and Jeremy P. Derrick. 2016. “Structures of Type IV Pilins from Thermus Thermophilus Demonstrate Similarities with Type II Secretion System Pseudopilins.” Journal of Structural Biology 196 (3): 375–84. 10.1016/j.jsb.2016.08.006.

Kaushik, Sharbani, Haoze He, and Ross E. Dalbey. 2022. “Bacterial Signal Peptides-Navigating the Journey of Proteins.” Frontiers in Physiology 13: 933153. 10.3389/fphys.2022.933153.

Korotkov, Konstantin V., and Wim G. J. Hol. 2008. “Structure of the GspK-GspI-GspJ Complex from the Enterotoxigenic Escherichia Coli Type 2 Secretion System.” Nature Structural & Molecular Biology 15 (5): 462–68. 10.1038/nsmb.1426.

Korotkov, Konstantin V., Maria Sandkvist, and Wim G. J. Hol. 2012. “The Type II Secretion System: Biogenesis, Molecular Architecture and Mechanism.” Nature Reviews. Microbiology 10 (5): 336–51. 10.1038/nrmicro2762.

Lam, Anita Y., Els Pardon, Konstantin V. Korotkov, Wim G. J. Hol, and Jan Steyaert. 2009. “Nanobody-Aided Structure Determination of the EpsI:EpsJ Pseudopilin Heterodimer from Vibrio Vulnificus.” Journal of Structural Biology 166 (1): 8–15. 10.1016/j.jsb.2008.11.008.

Makarova, Kira S., Eugene V. Koonin, and Sonja-Verena Albers. 2016. “Diversity and Evolution of Type IV Pili Systems in Archaea.” Frontiers in Microbiology 7 (May): 667. 10.3389/fmicb.2016.00667.

Mirdita, Milot, Konstantin Schütze, Yoshitaka Moriwaki, Lim Heo, Sergey Ovchinnikov, and Martin Steinegger. 2022. “ColabFold: Making Protein Folding Accessible to All.” Nature Methods 19 (6): 679–82. 10.1038/s41592-022-01488-1.

Nunn, D. N., and S. Lory. 1991. “Product of the Pseudomonas Aeruginosa Gene pilD Is a Prepilin Leader Peptidase.” Proceedings of the National Academy of Sciences of the United States of America 88 (8): 3281–85. 10.1073/pnas.88.8.3281.

Páez-Pérez, Edgar D., Araceli Hernández-Sánchez, Elvia Alfaro-Saldaña, and J. Viridiana García-Meza. 2025. “Simplified Method for Purifying Full-Length Major Type IV Pilins: PilA from Acidithiobacillus Thiooxidans.” MethodsX 15 (December): 103520. 10.1016/j.mex.2025.103520.

Parge, H. E., S. L. Bernstein, C. D. Deal, et al. 1990. “Biochemical Purification and Crystallographic Characterization of the Fiber-Forming Protein Pilin from Neisseria Gonorrhoeae.” The Journal of Biological Chemistry 265 (4): 2278–85.

Ramboarina, Stéphanie, Paula Fernandes, Peter Simpson, Gad Frankel, Michael Donnenberg, and Stephen Matthews. 2004. “Complete Resonance Assignments of Bundlin (BfpA) from the Bundle-Forming Pilus of Enteropathogenic Escherichia Coli.” Journal of Biomolecular NMR 29 (3): 427–28. 10.1023/B:JNMR.0000032511.89525.64.

Sheffield, Peter, Sarah Garrard, and Zygmunt Derewenda. 1999. “Overcoming Expression and Purification Problems of RhoGDI Using a Family of ‘Parallel’ Expression Vectors.” Protein Expression and Purification 15 (1): 34–39. 10.1006/prep.1998.1003.

Singh, Pradip Kumar, Janay Little, and Michael S. Donnenberg. 2022. “Landmark Discoveries and Recent Advances in Type IV Pilus Research.” Microbiology and Molecular Biology Reviews: MMBR 86 (3): e0007622. 10.1128/mmbr.00076-22.

Studier, F. William. 2005. “Protein Production by Auto-Induction in High-Density Shaking Cultures.” Protein Expression and Purification 41 (1): 207–34. 10.1016/j.pep.2005.01.016.

Teufel, Felix, José Juan Almagro Armenteros, Alexander Rosenberg Johansen, et al. 2022. “SignalP 6.0 Predicts All Five Types of Signal Peptides Using Protein Language Models.” Nature Biotechnology 40 (7): 1023–25. 10.1038/s41587-021-01156-3.

Winther-Larsen, Hanne C., Matthew Wolfgang, Steven Dunham, et al. 2005. “A Conserved Set of Pilin-like Molecules Controls Type IV Pilus Dynamics and Organelle-Associated Functions in Neisseria Gonorrhoeae.” Molecular Microbiology 56 (4): 903–17. 10.1111/j.1365-2958.2005.04591.x.

Yanez, Marissa E., Konstantin V. Korotkov, Jan Abendroth, and Wim G. J. Hol. 2008. “Structure of the Minor Pseudopilin EpsH from the Type 2 Secretion System of Vibrio Cholerae.” Journal of Molecular Biology 377 (1): 91–103. 10.1016/j.jmb.2007.08.041.

